# A Multiplex Droplet Digital PCR Assay for Quantification of HTLV-1c DNA Proviral Load and T-Cells from Blood and Respiratory Exudates Sampled in a Remote Setting

**DOI:** 10.1101/364570

**Authors:** David Yurick, Georges Khoury, Bridie Clemens, Liyen Loh, Hai Pham, Katherine Kedzierska, Lloyd Einsiedel, Damian Purcell

## Abstract

During human T-cell leukemia virus type-1 (HTLV-1) infection the frequency of cells harboring an integrated copy of viral cDNA, the proviral load (PVL), is the main risk factor for progression of HTLV-1-associated diseases. Accurate quantification of provirus by droplet digital PCR (ddPCR) is a powerful diagnostic tool with emerging uses for monitoring viral expression. Current ddPCR techniques quantify HTLV-1 PVL in terms of whole genomic cellular material, while the main target of HTLV-1 infection is the CD4^+^ and CD8^+^ T-cell. Our understanding of HTLV-1 proliferation and the amount of viral burden present in different compartments is limited. Recently a sensitive ddPCR assay was applied to quantifying T-cells by measuring loss of germline T-cell receptor genes as method of distinguishing non-T-cell from recombined T-cell DNA. In this study, we demonstrated and validated novel applications of the duplex ddPCR assay to quantify T-cells from various sources of human gDNA extracted from frozen material (PBMCs, bronchoalveolar lavage, and induced sputum) from a cohort of remote Indigenous Australians and then compared the T-cell measurements by ddPCR to the prevailing standard method of flow cytometry. The HTLV-1c PVL was then calculated in terms of extracted T-cell gDNA from various compartments. Because HTLV-1c preferentially infects CD4^+^ T-cells, and the amount of viral burden correlates with HTLV-1c disease pathogenesis, application of this ddPCR assay to accurately measure HTLV-1c-infected T-cells can be of greater importance for clinical diagnostics, prognostics as well as monitoring therapeutic applications.

## Introduction

Globally, HTLV-1 is estimated to infect around 20 million people who mostly reside in areas of high endemicity such as southwestern Japan, the Caribbean, South America, sub-Saharan Africa and the Mashhad district of Iran [1]. Recently, it was confirmed that a very high prevalence of HTLV-1 subtype C (HTLV-1c) infection occurs among Aboriginal adults in Central Australia, where prevalence rates exceed 40% in some remote communities [2]. Human T-cell leukemia virus type 1 (HTLV-1) is a lymphoproliferative and ultimately oncogenic retrovirus that primarily infects CD4^+^ T-cells [3] and is the causative agent of adult T-cell leukemia/lymphoma, HTLV-1-associated myelopathy/tropical spastic paraparesis [4, 5] and various other immune-mediated disorders [6-10]. In remote Australia, HTLV-1 infections are most significantly associated with bronchiectasis and multiple blood stream bacterial infections [2, 11, 12]. The HTLV-1 viral DNA burden is measured as the proviral load (PVL), which is the proportion of peripheral blood mononucleated cells (PBMCs) carrying an integrated copy of the HTLV-1 viral DNA. PVL correlates with the risk of disease development [13-17], however, levels of provirus can vary greatly between individuals, which complicates the prognostic use of this biomarker. Absolute quantification of the HTLV-1 PVL by ddPCR is a sensitive diagnostic tool with emerging applications for monitoring viral expression [18].

The main target for HTLV-1 infection, T-cells, are distinguished by the presence of a unique cell surface markers, such as CD3, CD4 and CD8, and their receptor for antigen termed the T-cell receptor (TCR) [19] (Figure 1). Most TCRs are composed of an alpha (α) and a beta chain (β) heterodimer, while a small proportion of T-cells that lacks TCRαβ chains expresses an alternative T-cell receptor, TCRγδ, with gamma (γ) and delta (δ) chains. The majority of T-cells undergo rearrangement of their TCRαβ through somatic rearrangement of multiple variable (V), diversity (D) and joining (J) gene segments at the DNA level [20]. V(D)J recombination occurs in developing lymphocytes during the early stages of T-cell maturation [21]. The first recombination event to occur is between one D and one J gene segment in the β chain of the TCR. This process could result in joining of the Dβ1 gene segment to any one of the six Jβ1 segments, or the Dβ2 gene segment to any one of the six Jβ2 segments. D-J recombination is followed by the joining of one Vβ-segment from an upstream region of the newly formed D-J complex, resulting in a rearranged V(D)J gene segment [20, 22]. All other gene segments between V and D segments are eventually deleted from the cell’s genome as a T-cell receptor excision circle (TREC) [23, 24]. The V(D)J transcript generated will incorporate the constant (C) region resulting in a Vβ-Dβ-Jβ-Cβ gene segment. Processing of the primary RNA adds a polyA tail after the Cβ and removes unwanted sequence between the V(D)J segment and the constant gene segment [25]. The levels of the different functional T-cells and proportions of their individual subtypes circulating in blood can vary significantly.

**Figure 1:**
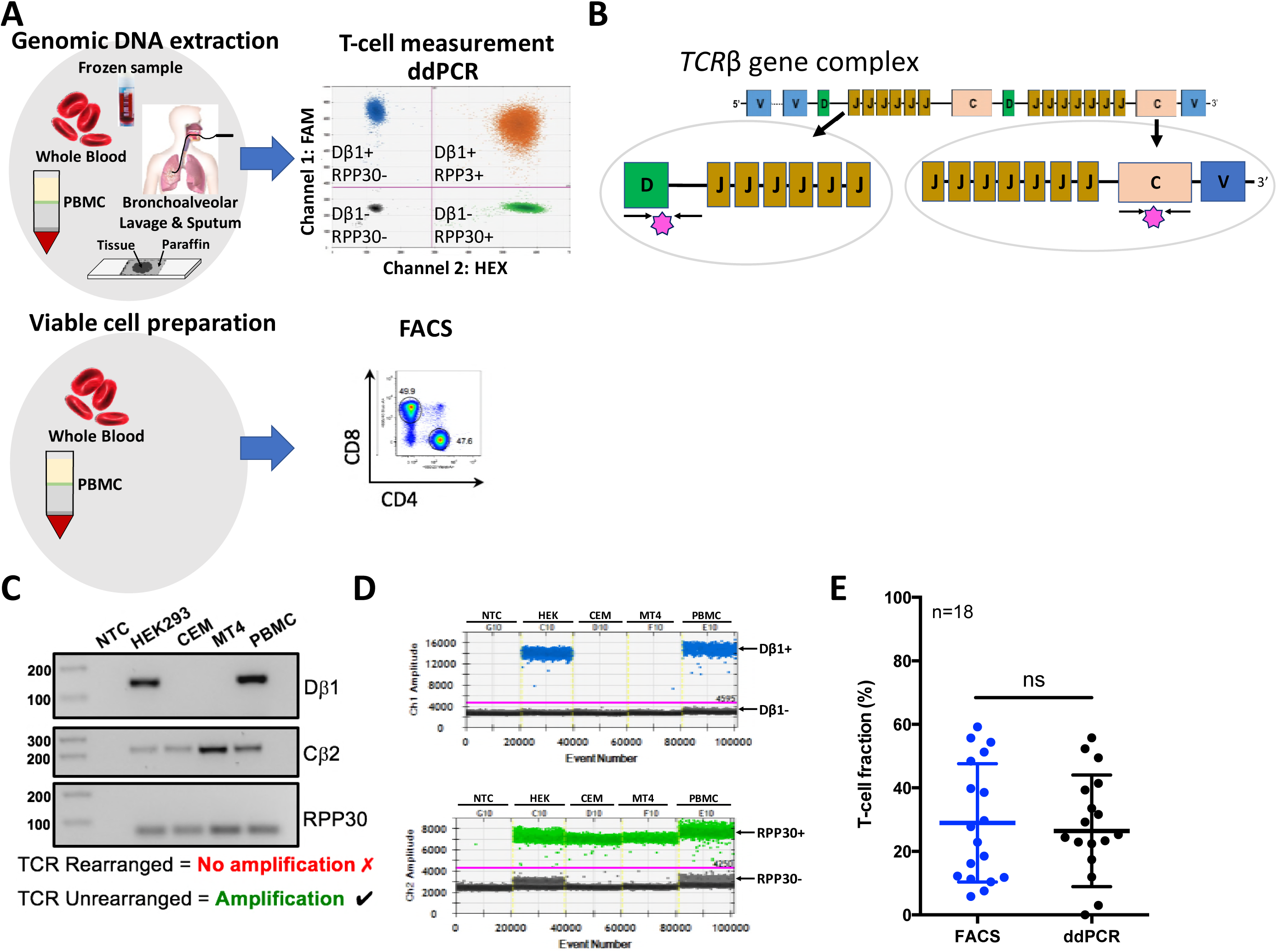
Validation of T-cell measurement by targeting the unrearranged T-cell receptor in comparison to flow cytometry. A)Study design and sample composition. Extracted genomic DNA from frozen blood, PBMCs, bronchoalveolar lavage and sputum samples obtained from remote Australian Indigenous HTLV-1c cohort was used to measure T-cells by a generic single duplex ddPCR assay. Viable cellular material isolated in whole blood and PBMCs from the same HTLV-1c cohort was used to measure T-cells by the gold standard method of flow cytometry. **B)** Schematic depiction of T-cell receptor α (TCRα) loci and the oligonucleotides (black arrows) and probes (pink star) used for detecting non T-cells (Diversity Dα - Joining jα) and all cells (constant region-2, Cβ). **C)** Validation of oligonucleotide specificity for detecting TCRβ rearrangement. Only cells that have not undergone TCR rearrangement present intact Dα-jα primer-binding regions and will result in a 143-base pair amplicons (noted Dα). The Cβ primers resulted in a 218-base pair amplicons since this region remains intact at the DNA level during VDJ recombination. The RPP30 primers resulted in a 62-base pair amplicons of all samples containing human gDNA. NTC, non-template control. **D)** A one-dimensional (1-D) ddPCR profile on Chi demonstrates the Dβ1 primer specificity to amplify samples containing non-T-cells or cells that have not undergone VDJ recombination (HEK and PBMC) (Dα+ blue droplets; Dα - black droplets); Ch2 1-D profile targeting the ubiquitous housekeeping gene, RPP30 (RPP30+ green droplets; RPP30-black droplets), which allows absolute quantification of total cells. Amplitude threshold is represented with a pink line. **E)** Comparison of T-cell quantification by FACS to ddPCR. Determined T-cell fractions of 18 healthy PBMC donors are plotted jointly for direct comparison of the two quantification methods. Bars indicate mean values with standard deviation (FACS: 29±18.6; ddPCR: 26±17.6) (Wilcoxon matched pairs test, p=0.6705, ns = non-significant).

Recently, a novel single duplex ddPCR assay was developed and validated for quantifying T-cells by measuring the loss of germline T-cell receptor loci, which resulted in accurate measurement of the T-cell population compared with the gold-standard method of flow cytometry [26]. The dynamic range of this technique makes certain that even low proportions of T-cells are accurately detected. In contrast to other techniques (flow cytometry, immunohistochemistry, real-time quantitative PCR), the digital design of ddPCR offers direct quantification and requires small amounts of DNA derived from fresh, frozen or fixed samples. This is particularly advantageous in a remote community setting where large distances and poor access to resources make it difficult to maintain cell viability of clinical samples, which often vary considerably in quantity and quality. Here, we describe a novel application of the recently introduced duplex ddPCR assay to quantify T-cells from various sources of human gDNA extracted from frozen material such as blood/PBMCs, bronchoalveolar lavage (BAL) and sputum samples obtained ethically from a remote Indigenous Australian HTLV-1 cohort.

## Results

### Quantification of T-Cells by Measuring the Unrearranged T-cell Receptor DNA

During early stages of T-cell maturation, rearrangements of Dβ1-Jβ1 intergenic sequences occur at both alleles, resulting in deletion of these sequences in nearly all peripheral T-cells [21]. In contrast, the TCRβ constant region-2 (Cβ2) remains intact during VDJ recombination. By measuring the loss of these specific TCRβ loci by ddPCR and normalizing against a stable reference gene, such as RPP30, enables a quantification of the number of T-cells in a clinical sample. On this basis, we designed a set of primer-probes that target the intact TCRβ gene region spanning across 143 base pairs of the Dβ1 exon and Jβ1 intron (Figure 1B). An additional primer-probe set were specifically designed to span 218 base pairs of the Cβ2 region and used as a positive control (Table 1). We validated our chosen Dβ1-Jβ1 target sequence for the detection of cells that had not undergone the VDJ recombination and thus were not capable of functioning as T cells using several different cell sources with varying T-cell composition. As expected, only cells that had not undergone T-cell rearrangement such as HEK293T and a subset of PBMCs comprising macrophages, monocytes, NK and B cells with intact primer binding regions resulted in a specific Dβ1-Jβ1 amplification (Figure 1C). On the other hand, T-cell lines MT4 and CEM that had clonally rearranged TCR genes failed to amplify the deleted TCR segment. All samples resulted in Cβ2 amplification since this region remains intact during VDJ recombination. Similarly, the results of a multiplexed ddPCR reaction confirmed that the amplification with Dβ/Jβ (CH1, FAM) is restricted to samples containing non-T cells that have not undergone VDJ recombination, while RPP30 reference gene (CH2, HEX) was detected for all samples (Figure 1D).

**Table 1:**
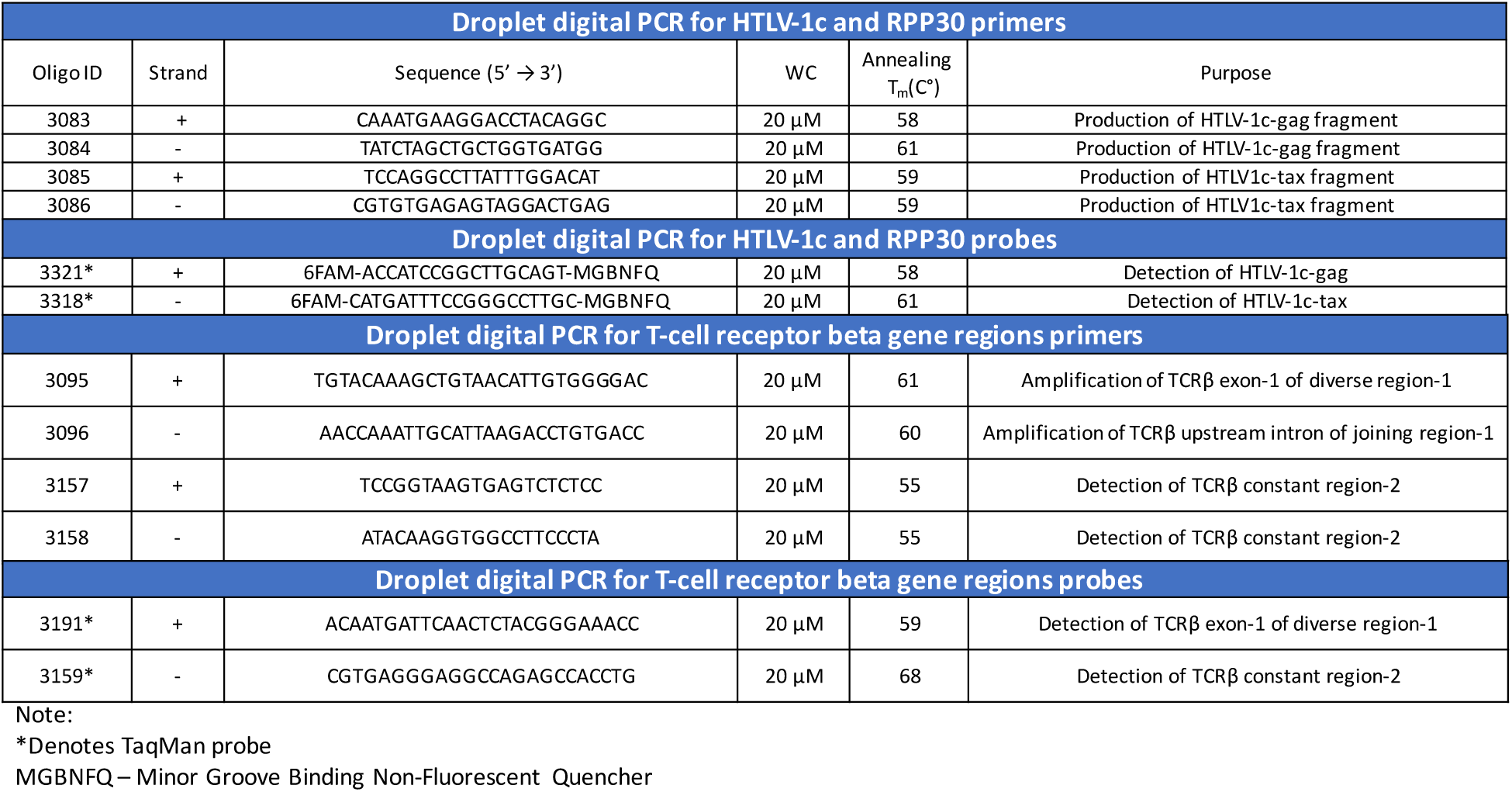
Primers and probe details used for ddPCR quantification of HTLV-1c and T-cells.

We validated this novel ddPCR assay against the gold standard flow cytometry method for T-cell measurement using CD3+ surface staining by comparing the ddPCR and the FACS determinations of the T-cell fraction from 18 healthy donor PBMC samples with varying levels of circulating T-cells (Figure 1E). No significant differences (p=0.6705, Wilcoxon matched-pairs test) in the frequency of CD3^+^ T cell fractions were detected between FACS (%29.0 ± 18.6) and ddPCR (%26.5 ± 17.6), confirming the specificity and accuracy in detecting unrearranged TCRβ and thus T-cells.

### High Accuracy and Dynamic Range of Detecting Unrearranged T-Cell Receptor DNA by ddPCR Technology

To evaluate the dynamic range of our unrearranged T-cell receptor (UTCR) assay, DNA isolated from non-T cells (HEK293T) were serially diluted into T-cell DNA (CEM cells) and evaluated by ddPCR with 4 replicates per sample. A comparison between the observed with the expected number of copies provided an estimation of the assay accuracy. The slope for the observed UTCR copy number (x: 0.808 ± 0.01) was significantly close to the expected UTCR copy number (y: 1.00 ± 0.0) (Figure Supplementary 1, R=0.9913, P<0.0001). The dynamic range of ddPCR from 1.56 to 10^5^ UTCR copies per well ensures sensitive and accurate detection of UTCR DNA, as indicated by the small 95% CIs. The ddPCR lower and upper limit of detection (LoD) for the UTCR assay was determined at 97.9 and 2×10^5^ copies per 10^6^ cells, respectively.

### Comparison of T-Cell Quantification in Sorted Cellular Populations Between ddPCR and Flow Cytometry Resulted in Positive Correlation

To further validate the assay utilized to calculate the T-cell population in gDNA of frozen samples, we compared measurements of FACS-sorted cell populations by flow cytometry and ddPCR. To perform this, we obtained PBMCs from healthy donors (n=6) and isolated various T-cell subsets (CD8^+^, CD4^+^ and *γ*δ^+^) and non-T-cell subsets (Natural killer [NK]-cells, monocytes and B-cells) by cell sorting based on expression of lineage-specific phenotypic markers as described in (Figure 2A). The purity checks resulted in ≥ 90% purity in most sorted populations, which reflects the overall homogeneity in each sorted group (Table 1). Two of the sorted healthy PBMC samples resulted in lower purity of *γ*δ^+^ T-cell populations, #5 (55.2%) and #6 (65.1%), reflecting possible down regulation of the *γ*δ TCR or photo bleaching during the sorting process. Lower purity was also noted in two of the sorted monocyte populations, #3 (78.5%) and #6 (77.7%), which could be attributed in part to the adherent nature of these larger cells that can result in adhesion to smaller cells despite attempts to gate out doublet cells. The percentage of T-cells measured by ddPCR in the T-cell populations was equivalent to the one determined by flow cytometry (Figure 2C, ns, p=0.7559, Wilcoxon matched-pairs signed rank test), which supports the functionality and the sensitivity of the UTCR assay to specifically detect and quantify T-cell populations. Moreover, the sorted non-T-cell population resulted in 0.0 to 8.6% total T-cells matching the percentage of purity observed by FACS (Table 1), which demonstrates the ability of the UTCR assay to accurately detect unrearranged TCRβ chain (Figure 2D, p <0.0001, r=0.9506, Pearson r test) and thus sorting efficiency and purity of T-cells from gDNA. The percentage of *γ*δ+ T-cells determined by FACS was 55.2 to 97%, while the ddPCR results ranged from 46.8 to 66% (Table 2). The 30% discrepancy between the flow cytometry and UTCR assay suggest that not all *γ*δ+ T-cells have fully rearranged TCRβ alleles.

**Figure 2A:**
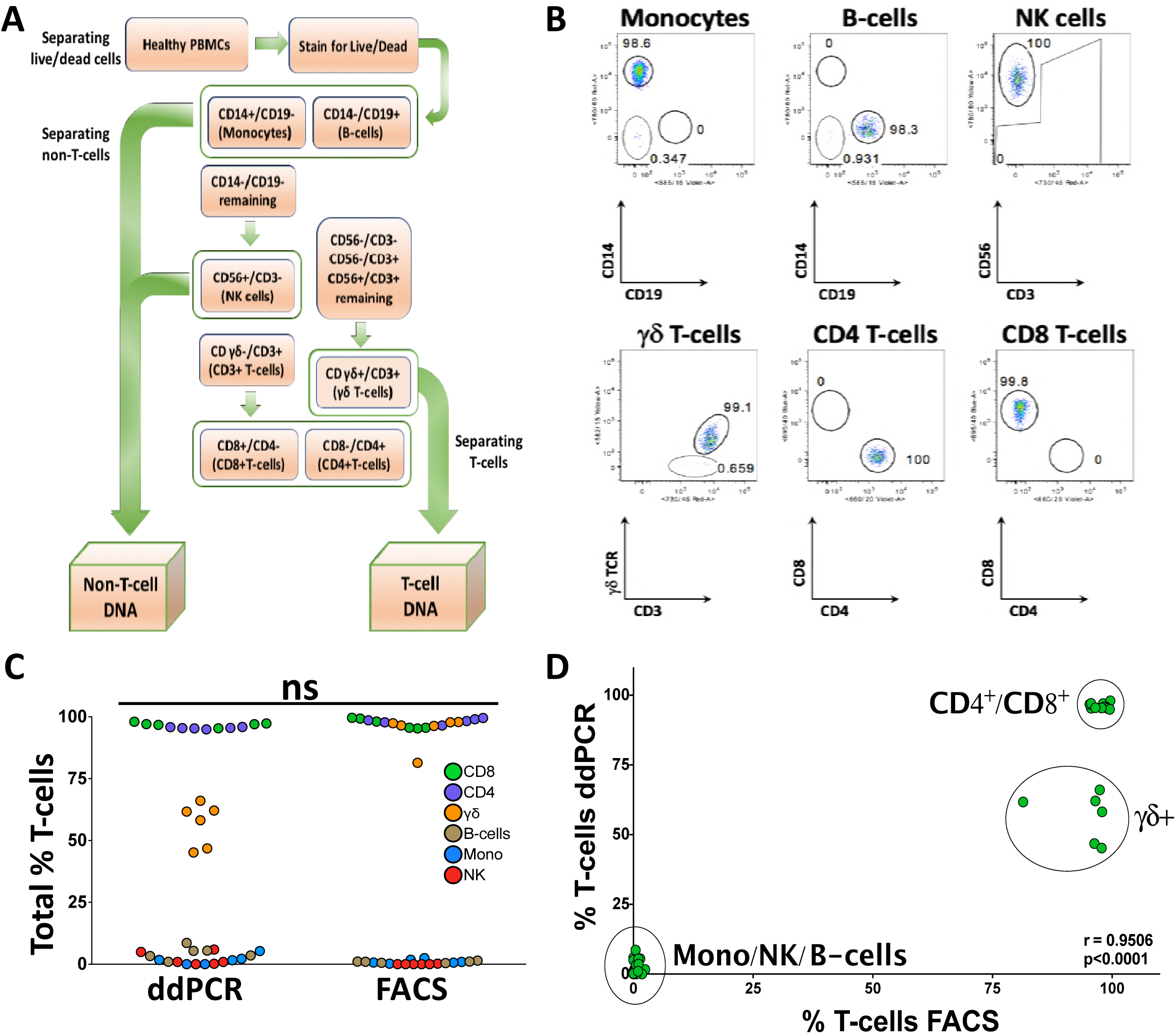
Comparison of T-cell quantification between ddPCR and flow cytometry in sorted cellular populations. **A)** Flowchart of FACS sorting strategy. PBMC samples from 6 healthy donors were sorted into non-T-cell (NK, monocyte and B-cells) and T-cell populations (CD8^+^, CD4^+^ and γδ), followed by DNA extraction. **B)** Purity checks of the various sorted cellular populations. **C)** Comparison of the total fraction of T-cells measured in each sorted population from healthy donors by ddPCR and FACS. Distribution of measured cell subsets was very similar, which did not result in a significant difference between the ddPCR and FACS assays (p=0.7559, Mann-Whitney). **D)** Correlation of ddPCR and FACS measured T-cells in sorted populations of T-cells and non-T-cells from healthy donors resulted in a positive correlation (p<0.0001, r=0.9506).

**Table 2:** Purity check of FACS-sorted cell populations and percentage T-cells measured by ddPCR.

| Purity Check |  |  |  |  |  |  |  |  |  |  |  |  |
| --- | --- | --- | --- | --- | --- | --- | --- | --- | --- | --- | --- | --- |
| Cell Type | % Purity of FACS-sorted Population |  |  |  |  |  | % T-cells in ddPCR measure |  |  |  |  |  |
| T-cells | PBMC1 | PBMC2 | PBMC3 | PBMC4 | PBMC5 | PBMC6 | PBMC1 | PBMC2 | PBMC3 | PBMC4 | PBMC5 | PBMC6 |
| CD8+ | 98.7 | 97.2 | 95.8 | 92.8 | 93.2 | 91.3 | 97.3 | 97.0 | 98.0 | 95.3 | 96.7 | 97.1 |
| CD4+ | 98.4 | 97.0 | 98.2 | 92.6 | 97.6 | 94.6 | 95.9 | 95.8 | 95.3 | 94.9 | 95.4 | 95.5 |
| γδ+ | 97.0 | 94.5 | 89.6 | 91.5 | 55.2 | 65.1 | 62.1 | 61.7 | 46.8 | 58.2 | 66.0 | 45.2 |
| Non-T-cells |  |  |  |  |  |  |  |  |  |  |  |  |
| NK | 99.5 | 99.9 | 99.0 | 95.1 | 94.4 | 95.3 | 0.9 | 0.0 | 0.9 | 5.9 | 4.9 | 0.2 |
| Monocyte | 92.9 | 89.5 | 78.5 | 93.9 | 92.5 | 77.7 | 1.6 | 5.3 | 0.0 | 1.7 | 2.2 | 0.0 |
| B-cell | 96.4 | 92.1 | 90.9 | 97.7 | 94.0 | 94.7 | 3.3 | 5.5 | 8.6 | 5.4 | 3.5 | 1.0 |

### Application of the UTCR Assay in a Remote Indigenous Australian HTLV-1c Cohort

We next examined the ability of the UTCR assay to quantify T-cells from various sources of HTLV-1 patient samples. To do this, we obtained frozen samples consisting of peripheral blood (n=29), induced sputum (n=6) and bronchoalveolar lavage (BAL) (n=3) from the Alice Springs Hospital-based Indigenous Australian HTLV-1 cohort, as well as blood samples from healthy donors of similar background (n=14). A summary of the participant characteristics and results is summarized in Supplementary Table 1 and Supplementary Table 2. The collection dates of the samples ranged from 21 January 2012 - 08 November 2016. The overall distribution of samples by gender was 22 males (57.9%) and 14 females (36.8%), with 2 unknown samples. The average age at time of sample collection was not significantly different between males (46.4 ± 2.9 years) and females (48.6 ± 2.3 years) (p=0.2718, unpaired t-test).

Given that HTLV-1 preferentially infects CD4^+^ T-cells, we hypothesized that the HTLV-1 PVL per T-cells would be higher in comparison with the PVL per genome since the latter includes all potential cellular targets of HTLV-1 infection and thus would dilute the PVL measurement. Collectively, the 38 samples resulted in a significant difference between the HTLV-1c PVL per genome and PVL per T-cell assays (Figure 3, p<0.0001, two-tailed Paired T-test), which indicates that the HTLV-1 PVL per T-cell assay quantifies a specific HTLV-1 targeted cellular population that could be relevant to the assessment of increased risk of HTLV-1 disease progression.

**Figure 3:**
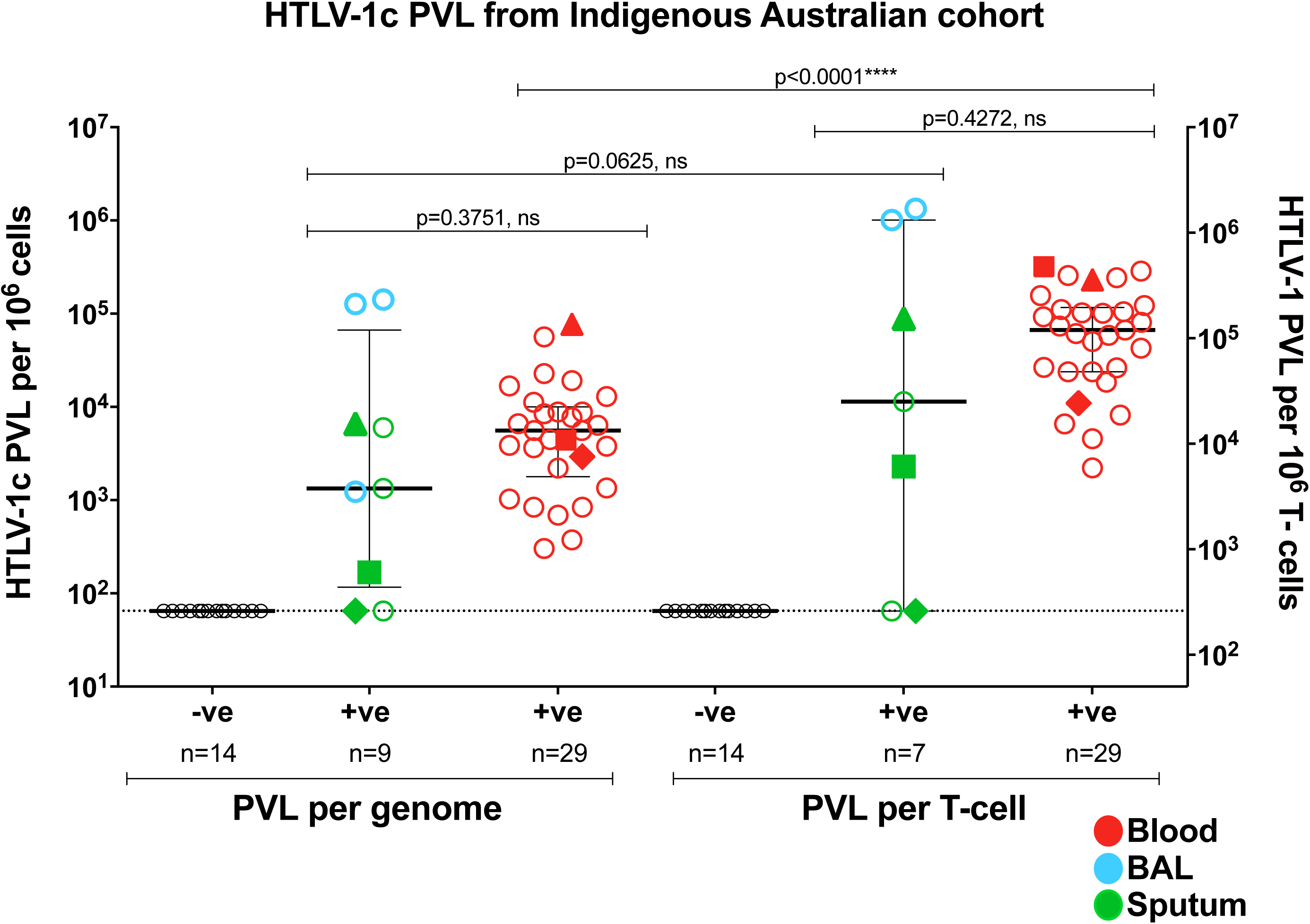
Distribution of HTLV-1c PVL measured in peripheral blood and various exudates from an Indigenous Australian cohort. HTLV-1c proviral load (PVL) per genome and PVL per T-cell were measured in HTLV-1c infected (+ve) peripheral blood (red), induced sputum (green) and bronchoalveolar lavage (BAL, blue) samples from remote Indigenous Australian cohort participants. PBMCs from healthy indigenous volunteers (-ve) were used as a negative control (open black circles). Box mid-line represents median value with interquartile range. Three subjects donated both blood and sputum samples designated by Δ, □ and *****. Isolated gDNA from one BAL and one sputum sample was insufficient for PVL per T-cell assay.

Next, we investigated whether the HTLV-1 PVL was consistent between these sources of infected blood and various inflammatory exudates (Figure Supplementary 3). The median and interquartile range (IQR) for HTLV-1c PVL per genome and PVL per T-cell in peripheral blood was 5.6×10^3^ copies (IQR, 1.8×10^3^, 1.0×10^4^) (per 10^6^ cells) and 6.7×10^4^ copies (IQR, 2.4×10^4^, 1.2×10^5^) (per 10^6^ T-cells), respectively. The median and IQR for HTLV-1c PVL per genome and PVL per T-cell in BAL was 1.3×10^5^ copies (IQR, 1.2×10^3^, 1.4×10^5^) (per 10^6^ cells) and 1.2×10^6^ copies (IQR, 1.0×10^6^, 1.3×10^6^) (per 10^6^ T-cells), and 754.0 copies (IQR, 64.8, 6.1×10^3^) (per 10^6^ cells) and 2.3×10^4^copies (IQR, 97.8, 5.0×10^4^) (per 10^6^ T-cells) respectively in the induced sputum.

We observed a significantly higher mean ± SEM of HTLV-1c PVL per genome in blood (1.1×10^4^ ± 3.1×10^3^ copies/per 10^6^ cells) compared with induced sputum (2.4×10^3^ ± 1.3×10^3^ copies/per 10^6^ cells; p=0.0388, unpaired t-test). We also observed a significantly higher mean ± SEM HTLV-1c PVL per T-cell in blood (9.4×10^4^ ± 1.7×10^4^ copies/per 10^6^ T-cells) compared with induced sputum (2.0×10^4^ ± 1.7×10^4^ copies/per 10^6^ T-cells; p=0.0133, unpaired t-test). Overall, the mean PVL per genome and PVL per T-cell in blood was approximately 4 -and 5-times greater than in sputum, respectively.

We also observed the mean ± SEM HTLV-1c PVL per genome was higher in BAL (9.0×10^4^ ± 4.5×10^4^ copies/per 10^6^ cells) compared with that of blood samples, although this result was not significant. However, the mean ± SEM HTLV-1c PVL per T-cell in BAL (1.2×10^6^ ± 1.6×10^5^ copies/per 10^6^ T-cells) was significantly higher compared with that of blood samples (p=0.0043, unpaired t-test). The mean PVL per genome and PVL per T-cell in BAL samples was approximately 9- and 13-times higher than blood, respectively.

Finally, we observed that the mean ± SEM HTLV-1c PVL per genome and PVL per T-cell in induced sputum (2.4×10^3^ ± 1.3×10^3^ copies/per 10^6^ cells) and (2.0×10^4^ ± 1.7×10^4^ copies/per 10^6^ T-cells), respectively, was lower compared with that of BAL samples, although neither result reached statistical significance. The PVL per genome and PVL per T-cell in BAL samples was 38- and 57-times higher than induced sputum samples, respectively.

## Discussion

Previous studies have shown that quantitative PCR (qPCR) and ddPCR are both capable of distinguishing clinically significant differences in T-cell proportions and perform similarly to FACS [27]. However, ddPCR technology results in a high-throughput digital PCR with several advantages over qPCR [28, 29]. Unlike qPCR, ddPCR provides an absolute count of target copies independent of an extrapolation from a standard curve, which greatly reduces variability between assays and difficulty in measuring PVL, particularly from samples with low numbers of cells [30, 31]. Direct measurement of target DNA is optimal for viral load analysis, and when combined with the massive sample partitioning afforded by ddPCR, a greater precision and reliability can be achieved [32].

We have demonstrated and validated a novel application of the ddPCR assay to accurately measure T-cells in HTLV-1-infected peripheral blood and inflammatory exudates. Specifically, we provided evidence from a remote Indigenous Australian HTLV-1c cohort that the viral burden varies between compartments. Collectively, we found a significant difference between HTLV-1 PVL per genome and PVL per T-cell, indicating that the HTLV-1 PVL per T-cell assay quantifies a specific HTLV-1 targeted cellular population that could be relevant to the assessment of increased risk of HTLV-1 disease progression. A higher HTLV-1 proviral burden resides in cells extracted from BAL samples compared with peripheral blood, and suggests differences in the location of the HTLV-1 inflammatory response. In fact, measurement of specific compartments such as the lungs may be a better indicator for risk of disease progression, specifically HTLV-1c-associated respiratory diseases such as bronchiectasis [11].

Given the difficulty in collecting clinical material from remote community setting, there are several limitations in this study such as the limited number of subjects who provided pulmonary secretions. Further work is necessary to confirm our findings in larger studies in central Australia. It is also critical to compare PVL from different compartments in the same individual given that wide variability in peripheral blood PVL exists between individuals, which could complicate comparisons between groups and the use of PVL as a prognostic tool. In addition, low cell numbers are more likely to explain why we measured such low PVL in the induced sputum. The range of cells and T-cells for sputum samples was 264.5 - 640.5 and 13.0 - 74.5, respectively. While the sputum cell numbers were low compared to BAL samples, the BAL was only collected during procedures under the setting of intensive care making BAL samples unsuitable for monitoring of HTLV-1 involvement in lung disease. A further limitation results from the lack of clinical history of these subjects. Without this information, potential complications such as pulmonary disease or infective exacerbation during sample collection could influence our results.

In conclusion, our data supports the application of the ddPCR assay to count T-cells from DNA specimens from various compartments, and has potential clinical and diagnostic applications in the sharply focused longitudinal monitoring of HTLV-1 PVL and risk assessment of HTLV-1-associated inflammatory diseases. Furthermore, this assay has translational applications in the validation of cell purity following isolation of CD4^+^, CD8^+^, *γ*δ^+^ T-cells, as well as B-cells, NK cells and monocytes. In order to fully explore the applications of this UTCR assay, it will be essential to conduct larger HTLV-1 case-controlled studies and experimentally address how the viral burden in specific compartments correlates with HTLV-1-associated disease pathogenesis.

### Materials and Methods

#### Primary Cells

Whole blood samples from HTLV-1c patients were collected from adult subjects (age ≥ 18 years) who were recruited >48h after admission to Alice Springs Hospital, Northern Territory, central Australia, between 21 January 2012 - 08 November 2016. With ethics approval and patient consent in primary language, frozen specimens consisting of 29 peripheral blood, 6 induced sputum and 3 bronchoalveolar lavage (BAL) samples from the remote Indigenous Australian cohort were sent to The Peter Doherty Institute for Infection and Immunity at The University of Melbourne. Also, 14 healthy subjects from similar background were included as negative controls. gDNA was extracted using GenElute^™^ Blood Genomic DNA Kit (Sigma-Aldrich) according to manufacturer’s instructions and eluted in EB buffer (Sigma-Aldrich) or RNA-free water. To ensure efficient gDNA extraction from sputum and BAL, samples were supplemented with carrier DNA and treated for 3 h at 55°C with lysis buffer and proteinase K (200 μg). Purity of the isolated DNA A260/280 ratio was measured by UV spectrophotometry (Nanodrop Technologies, Wilmington, CA).

#### HTLV-1 Serologic and Molecular Studies

HTLV-1 serostatus was based on the detection of specific anti-HTLV-1 antibodies in serum by enzyme immunoassay (EIA) (Murex HTLVI+II; DiaSorin, Saluggia, Italy) and the Serodia^®^HTLV-I particle agglutination assay (Fujirebio, Tokyo, Japan) performed by the National Serological Reference Laboratory, Melbourne, Australia.

#### ddPCR Limit of Detection of HTLV-1c gag and tax

To evaluate the dynamic range and accuracy of quantifying HTLV-1 gene regions by ddPCR, a 1:5 serial dilution of plasmids containing HTLV-1c viral targets (pCRII-HTLV1c-*gag* and pCRII-HTLV1c-*tax*) were used to determine the lower and upper LoD. The standard curve was performed in duplicate as independent experiments, resulting in partitioning of approximately 40,000 droplets. Where the data points stray from linearity represents the lower and upper LoD.

#### ddPCR Limit of Detection of Non-T-cells

To evaluate the dynamic range and accuracy of quantifying T-cells by ddPCR, a 1:5 serial dilution of gDNA isolated from non-T-cells (HEK 293T) and T-cells (CEM) (each 6×10^6^ cells/ml) were used to determine the lower and upper LoD in measuring the number of unrearranged TCRβ gene regions. CEM T-cells were added to each well to maintain normalized levels of gDNA throughout the assay. The intact TCRβ gene region spanning across the Dβ1 and Jβ1 region was measured in duplicate for each sample on 3 separate occasions for n=3.

#### ddPCR HTLV-1 PVL Measurements

To quantify the PVL accurately, primers (900nM) and FAM-conjugated hydrolysis probes (250nM) specific to a conserved HTLV-1c-gag or -tax were developed (Table 1). Probes targeting the provirus were labeled with FAM (Applied Biosystems), whereas the probe directed at the reference gene *RPP30* (Ribonuclease P/MRP subunit P30, dHsaCPE5038241, Bio-Rad) was labeled with HEX. All primers and probes were designed for ddPCR and cross-checked with binding sites against the human genome to ensure target specificity of the generated primer pairs (Primer-BLAST, NCBI). A temperature optimization gradient ddPCR assay was performed to determine the optimal annealing temperature of primers targeting HTLV-1 *gag* and *tax* (data not shown). ddPCR was performed using ddPCR Supermix for probes (no dUTP, Bio-Rad Laboratories, Hercules, CA) in 22 μl with 50-100 ng of gDNA. Following droplet generation (15,000-18,000 on average) using a QX-200 droplet generator, droplets were then transferred to a 96-well plate (Eppendorf, Hauppauge, NY), heat-sealed with pierceable sealing foil sheets (ThermoFisher Scientific, West Palm Beach, FL), and amplified using a C1000 Touch^™^ thermocycler (Bio-Rad) with a 105°C heated lid. Cycle parameters were as follows: enzymatic activation for 10 minutes at 95°C; 40 cycles of (denaturation for 30 seconds at 94°C, annealing and extension for 1 minute at 58°C); enzymatic deactivation for 10 minutes at 98°C; and infinite hold at 10°C. All cycling steps utilized a ramp rate of 2°C/sec. Droplets were analyzed with a QX200 droplet reader (Bio-Rad) using a two-channel setting to detect FAM and HEX. The positive droplets were designated based on the no template controls (NTC) and FMO controls (HTLV-1(-)/RPP30(+); HTLV-1(+)/RPP30(-) and HTLV-1(+)/RPP30(+)) using gDNA extracted from healthy donors, HTLV-1c tax plasmid (pcRII-tax) and MT4 gDNA, which were included in each run. While our primers are specific for HTLV-1c, they work efficiently in detecting HTLV-1a from MT4 cell line [18].

#### ddPCR T-Cell Measurements

Methods to quantify T-cells accurately using the duplex ddPCR assays have been previously described by Zoutman et al., 2017 [26]. However, different primers and probe were utilized in this study (Table 1). Probes directed at the intact TCRβ gene region, which represents a cell that has not undergone VDJ recombination and spanning across 143 base pairs of the Dβ1 - Jβ1 region were labeled with FAM, whereas probes directed at the internal reference gene *RPP30* were labeled with HEX to quantify the total number of cells (Table 1). Additional primers and probe were specifically designed to span 218 base pairs of the TCRβ constant region-2 (Cβ) and used as a positive control (Table 1). The final concentrations of each primer and probe used in the ddPCR reaction were 900nM and 250nM, respectively. A temperature optimization gradient assay was performed to determine the optimal annealing temperature of primers targeting TCRβ gene regions (data not shown). ddPCR was performed as previously described, but the cycle parameters were as follows: enzymatic activation for 10 minutes at 95°C; 50 cycles of (denaturation for 30 seconds at 94°C, annealing and extension for 1 minute at 60°C); enzymatic deactivation for 10 minutes at 98°C; and infinite hold at 10°C.

#### ddPCR HTLV-1 PVL Data Analysis

QuantaSoft software version 1.7.4 (Bio-Rad) was used to quantify and normalize the copies/μl of each target per well. To address the HTLV-1-infected samples, which might be at or below the LoD, calculation of proviral copy number was normalized to the lower LoD of the PVL assay (65 copies per 10^6^ cells). Amplitude fluorescence thresholds were manually determined according to the negative controls (non-template control and DNA from healthy PBMCs), which had been included in each run. Droplet positivity was measured by fluorescence intensity above a minimum amplitude threshold. All samples were run in duplicate, and the HTLV-1 PVL was determined as the mean of the two measurements. The HTLV-1 PVL per genome was calculated based on the concentration of HTLV-1 target gene, either *gag* or tax, and expressed as proviral copies per μl, and divided by the copies of RPP30 diploid genome. The quotient is then multiplied by a chosen unit of cells designated as 1 × 10^6^ cells.

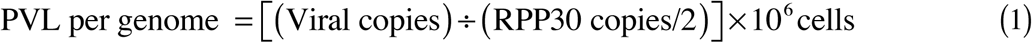

#### ddPCR T-cell Data Analysis

Quantification and normalization of number of T-cells was previously described [26]. Briefly, to address the HTLV-1c-infected samples, which might be at or below the LoD, calculation of the number of T-cells in each sample was normalized to the lower LoD of the UTCR assay (98 copies per 10^6^ T-cells). All samples were run in duplicate to quantify the absolute mean number of intact Dβ/Jβ-regions, or non-T-cells, which represents a cell that has not undergone VDJ recombination. As previously described by Zoutman et al., the total number of non-T-cells is quantified absolutely by ddPCR and then subtracted from the total number of cells to arrive at the total T-cell fraction. From this, the HTLV-1c PVL per T-cell was calculated based on the corresponding HTLV-1 PVL per genome values targeting *gag* or *tax*, and defined as the HTLV-1 proviral copies per 10^6^ T-cells. If the PVL per genome is derived from total genomic material, and the proportion of T-cells is calculated by subtraction from the proportion of non-T-cells, the contribution of T-cells to the PVL is calculated in the following manner:

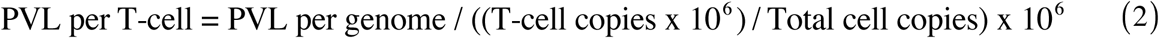

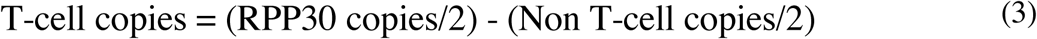

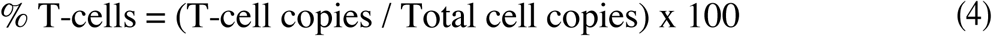

#### Flow Cytometry

Flow cytometry was performed on frozen PBMCs isolated from buffy coats (Australian Red Cross Blood Service, West Melbourne, Australia). Cryopreserved cells were rapidly thawed at 37°C, added dropwise to thawing media containing fresh cRPMI (Roswell Park Memorial Institute 1640 medium (RPMI; Gibco Invitrogen Cell Culture, Grand Island, NY, USA) with 10% fetal calf serum (FCS; Bovogen Biologicals, East Keilor, VIC, Australia), 2mM L-glutamine, 1mM sodium pyruvate, 100 μM MEM non-essential amino acids, 5mM HEPES buffer (all Gibco), 55 μM 2-mercaptoethanol (Invitrogen Corporation, Carlsbad, CA, USA), 100 U/ml penicillin and 100 U/ml streptomycin (both Gibco) and benzonase (50U/ml) (Novagen, ED Millipore Corporation, Billerica, MA, USA). Cells were then centrifuged at room temp for 6 min at 500 x g, counted and resuspended in PBS, and then stained with Live/Dead-Aqua (Molecular Probes for Life Technologies) to exclude potential autofluorescence from dead cells. Cells were then washed twice with PBS and stained with a combination of anti-CD3 Alexa Fluor 700 (UCHT1), anti-CD4 BV650 (SK3), anti-CD8 PerCPCy5.5 (SK1), anti-CD14 APC-H7 (MΦP9), anti-CD56 PE-Cy7 (NCAM16.2), anti-TCR-γδ-1 PE (11F2) and anti-CD19 BV711 (SJ25C1) (all BD Biosciences) (PBS with 0.1% Bovine Serum Albumin, Gibco for Life Technologies). After washing twice with sort buffer, cells were resuspended and passed through a 70μm sieve and acquired by Fluorescence-activated cell sorting (FACS; BD FACS Aria Fusion, BD Immunocytometry Systems, San Jose, CA, USA) to isolate live populations of non-T-cells (NK, B-cells, monocytes), T-cells (CD8^+^, CD4^+^, Ɣδ^+^). The flow gating strategy to sort non-T-cell and T-cell populations was as follows: live non-T-cell populations of B-cells (CD19+CD14^-^), Monocytes (CD14+CD19^-^), NK cells (CD14^-^CD19^-^CD56+CD3^-^); and then live T-cell populations of γδ^+^ T-cells (CD14^-^CD19^-^CD3^+^TCRγδ^+^), CD4^+^ T-cells (CD14^-^CD19^-^ CD3^+^TCRγδ^-^CD4^+^) and CD8^+^ T-cells (CD14^-^CD19^-^CD3^+^TCRγδ^-^CD8^+^) (Figure 2A). After sorting the samples into respective populations, a purity check for each population was subsequently performed. Gates were carefully chosen to reduce the selection of unspecific cellular populations. (Figure 2B). Data were analyzed with FlowJo version 9.7.6 (Tree Start) software.

#### Statistical Analysis

GraphPad Prism version 6 (GraphPad Software, La Jolla, CA) software was used for statistical analysis. To evaluate linear association in the fraction of T-cells measured between ddPCR and flow cytometry, linear regression and standard Pearson *r* tests were performed. T-cell quantification data from healthy and HTLV-1c-infected cohort samples were depicted as dot plots and tested for differences in median counts by Kruskal-Wallis testing with a confidence interval of 95%. Mann-Whitney was used to compare unpaired samples, and Paired T test was used to compare paired specimens (blood, BAL and sputum). P < 0.05 was considered significant.

## Supplemental Material

Supplemental material for this article may be found online.

## Acknowledgements

We would sincerely like to thank all the remote Indigenous Australian community members who participated in this study. We also would like to thank members of the scientific community who generously shared reagents critical to this work. We acknowledge Kim Wilson of the National Reference Laboratory of Melbourne, Australia, and gratefully acknowledge the support of the Pathology Department at Alice Springs Hospital. We would also like to thank the DMI Flow Facility staff for their advice and generous assistance during the sorting experiments.

The study was reviewed and approved by the Central Australian Human Research Ethics Committee. All patients were informed in first language and gave written informed consent in accordance with the National Health and Medical Research Council of Australia. (HREC-14-249). The datasets used and/or analyzed during the current study relates to Indigenous Australians and cannot be accessed without appropriate ethics approval from the Central Australian Human Research Ethics Committee for researchers who meet the criteria for access to confidential data.

The authors declare that they have no competing interests.

This study was supported by the National Health and Medical Research Council of Australia (NHMRC) program grant #1052979 to DP and program grant #1071916 to KK. KK is a NHMRC Senior Research Level B Fellow (#1102792) and BC is a NHMRC Peter Doherty Fellow.

---

**Figure Supplementary 1:**
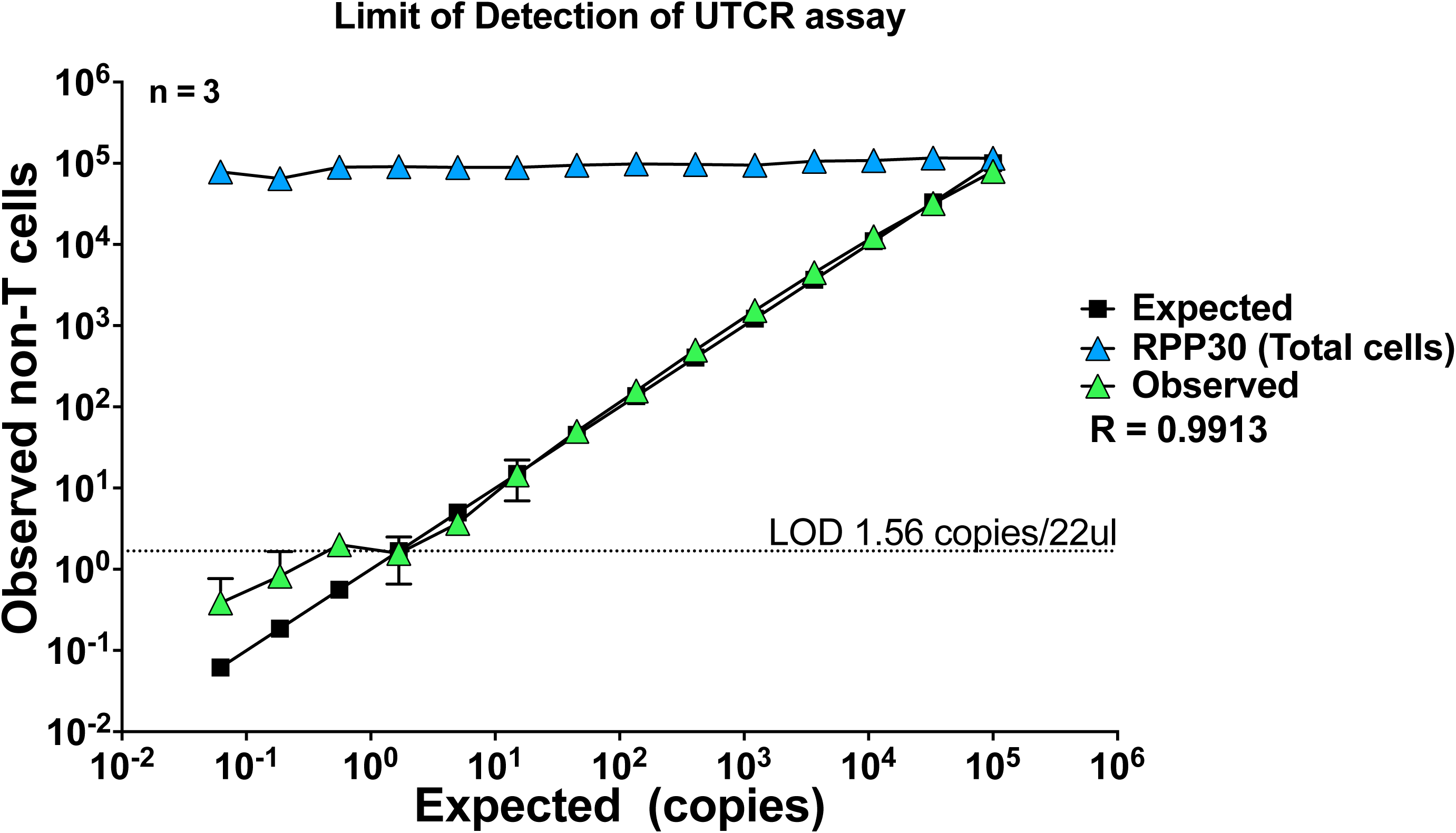
ddPCR limit of detection of UTCR assay. A 1:5 serial dilution of gDNA from HEK293T and CEM cells was performed to determine the limit of detection (LoD) of the UTCR assay. Data shown are the mean values of 3 independent measurements each conducted in duplicate (n=3). Comparison of the observed number of copies from each target (y-axis) with the expected number of copies (x-axis) provides an estimation of the assay accuracy. The dilution series strays from linearity at 1.56 copies per 22ul well. The ddPCR lower and upper LoD for the UTCR assay was determined at 97.9 and 2×10^6^ copies per 10^6^ cells, respectively.

**Figure Supplementary 2:**
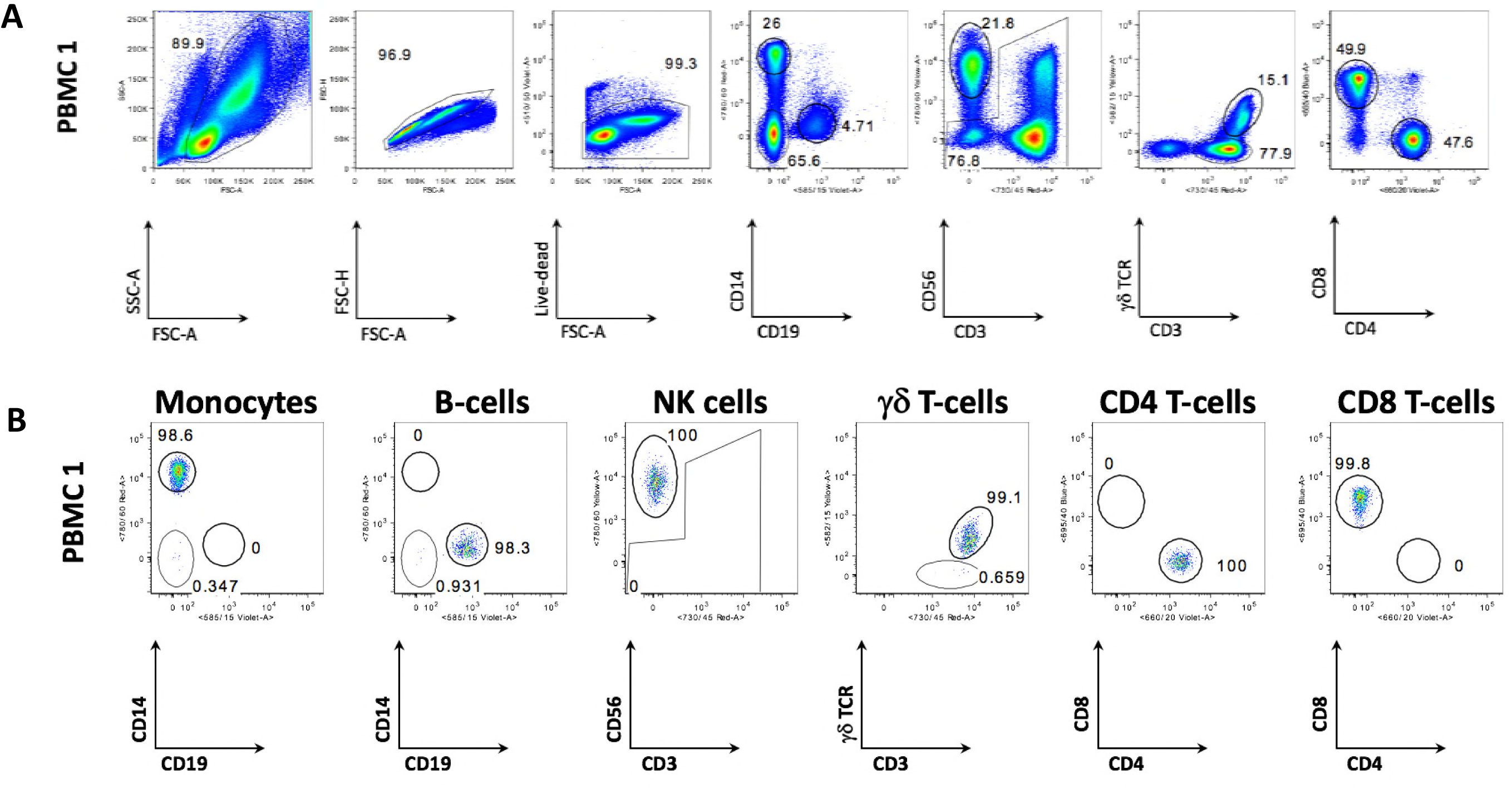
Sequential gating to identify specific leukocyte subsets. **A)**Gating strategy into various T-cell subsets (CD8^+^, CD4^+^ and ɣδ^+^ cells) and non-T-cell populations (NK, Monocytes and B-cells). B) Purity check of sorted populations and percentages of cells present in sorted samples.

**Table Supplementary 1:** Clinical characteristics and HTLV-1 proviral load (PVL) of 29 indigenous adult blood donors from remote Central Australia.

**Figure S2**
| Collection Date | Age | Sex | PVL per Genome<br>(copies per 10<sup>6</sup> cells) | PVL per T-cell<br>(copies per 10<sup>6</sup> T-cells) | Sample Type |
| --- | --- | --- | --- | --- | --- |
| 21-Nov-12 | 48 | M | 8.81E+03 | 5.86E+04 | Blood |
| 29-Nov-12 | 55 | F | 3.03E+02 | 4.26E+04 | Blood |
| 20-Nov-12 | 43 | F | 6.92E+02 | 6.58E+03 | Blood |
| 9-Jul-13 | 68 | M | 3.79E+03 | 6.67E+04 | Blood |
| 28-Feb-13 | 53 | F | 1.12E+04 | 9.26E+04 | Blood |
| 1-Mar-13 | 58 | M | 8.39E+02 | 1.84E+04 | Blood |
| 27-Feb-13 | 51 | F | 8.84E+03 | 8.07E+04 | Blood |
| 20-Aug-13 | 40 | M | 8.42E+02 | 8.19E+03 | Blood |
| 3-Oct-13 | 67 | M | 4.45E+03 | 5.02E+04 | Blood |
| 16-Oct-13 | 44 | M | 6.60E+03 | 2.57E+05 | Blood |
| 19-Nov-13 | 48 | M | 6.35E+03 | 1.11E+05 | Blood |
| 25-Nov-13 | 60 | M | 5.57E+03 | 1.22E+05 | Blood |
| 21-Feb-12 | 51 | F | 7.70E+03 | 6.13E+04 | Blood |
| 3-Feb-12 | 48 | F | 3.74E+02 | 2.22E+03 | Blood |
| 22-Nov-12 | 53 | M | 1.35E+03 | 2.63E+04 | Blood |
| 30-Sep-16 | 51 | F | 1.91E+04 | 7.35E+04 | Blood |
| 26-Oct-16 | 23 | M | 8.43E+03 | 1.01E+05 | Blood |
| 12-Jun-16 | 51 | F | 3.87E+03 | 2.38E+04 | Blood |
| 10-Jun-16 | 48 | M | 1.29E+04 | 1.56E+05 | Blood |
| 27-Feb-13 | 37 | M | 3.65E+03 | 1.05E+05 | Blood |
| 13-Feb-13 | 33 | M | 5.58E+03 | 2.38E+04 | Blood |
| 29-Aug-12 | 33 | F | 1.68E+04 | 2.43E+05 | Blood |
| 21-Jan-12 | 48 | M | 1.03E+03 | 4.55E+03 | Blood |
| 21-Nov-12 | 65 | F | 5.62E+04 | 2.87E+05 | Blood |
| 24-Sep-13 | 56 | F | 2.28E+04 | 1.02E+05 | Blood |
| 8-Nov-13 | 46 | M | 2.93E+03 | 1.10E+04 | Blood |
| 11-Jun-12 | 33 | F | 7.72E+04 | 2.32E+05 | Blood |
| 21-Oct-13 | 44 | M | 4.42E+03 | 3.19E+05 | Blood |
| 6-Nov-12 | 43 | M | 2.21E+03 | 2.66E+04 | Blood |

**Table Supplementary 2:** Detailed summary of 9 inflammatory exudate donors from remote Central Australian Indigenous HTLV-1c cohort.

| Collection Date | Age | Sex | PVL pergenome<br>(copies per 10<sup>6</sup> cells) | PVL per T-cell<br>(copies per 10<sup>6</sup> T-cells) | Sample Type |
| --- | --- | --- | --- | --- | --- |
| Unknown | Unknown | M | 5.98E+03 | 6.48E+01 | Sputum |
| 30-Sep-16 | 51 | F | 6.48E+01 | 6.48E+01 | Sputum |
| 26-Oct-16 | 23 | M | 6.62E+03 | 8.86E+04 | Sputum |
| 8-Nov-16 | 67 | M | 6.48E+01 | 6.48E+01 | Sputum |
| Unknown | 74 | M | 1.34E+03 | 1.14E+04 | Sputum |
| 12-Jun-16 | 51 | F | 6.48E+01 | 6.48E+01 | Sputum |
| 7-Nov-13 | 46 | M | 5.27E+03 | N/A | BAL RUL |
| 21-Dec-15 | 32 | M | 1.27E+05 | 1.01E+06 | BAL RUL |
| 21-Dec-15 | 32 | M | 1.42E+05 | 1.33E+06 | BAL LLL |

**Figure Supplementary 3:**
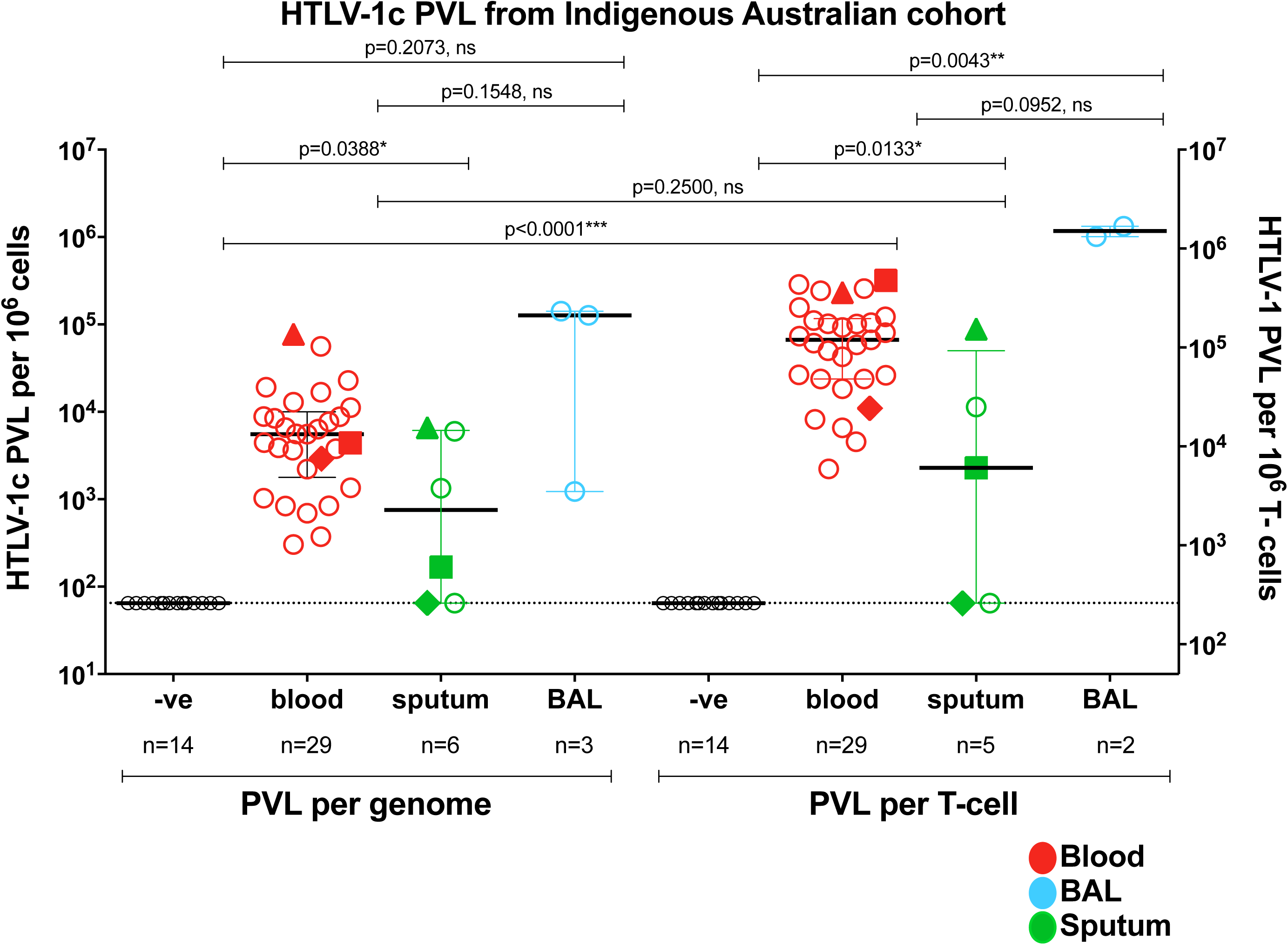
Relative distribution of HTLV-1c PVL measured in blood and inflammatory exudates from a remote Indigenous Australian cohort. Distribution of HTLV-1c proviral load (PVL) per genome and PVL per T-cell within peripheral blood (red), induced sputum (green) and bronchoalveolar lavage (BAL, blue) samples. Three subjects donated both blood and sputum samples designated by Δ, □ and ⋄. PBMCs from healthy indigenous volunteers (-ve) were used as a negative control (open black circles). Line represents median value with interquartile range. Isolated gDNA from one BAL and one sputum sample was insufficient for PVL per T-cell assay.

